# The influence of landscape and environmental factors on ranavirus epidemiology in amphibian assemblages

**DOI:** 10.1101/167395

**Authors:** Brian J. Tornabene, Andrew R. Blaustein, Cheryl J. Briggs, Dana M. Calhoun, Pieter T. J. Johnson, Travis McDevitt-Galles, Jason R. Rohr, Jason T. Hoverman

## Abstract

**Aim:** To quantify the influence of a suite of landscape, abiotic, biotic, and host-level variables on ranavirus disease dynamics in amphibian assemblages at two biological levels (site and host-level).

**Location:** Wetlands within the East Bay region of California, USA.

**Methods:** We used competing models, multimodel inference, and variance partitioning to examine the influence of 16 landscape and environmental factors on patterns in site-level ranavirus presence and host-level ranavirus infection in 76 wetlands and 1,377 amphibian hosts representing five species.

**Results:** The landscape factor explained more variation than any other factors in site-level ranavirus presence, but biotic and host-level factors explained more variation in host-level ranavirus infection. At both the site- and host-level, the probability of ranavirus presence correlated negatively with distance to nearest ranavirus-positive wetland. At the site-level, ranavirus presence was associated positively with taxonomic richness. However, infection prevalence within the amphibian population correlated negatively with vertebrate richness. Finally, amphibian host species differed in their likelihood of ranavirus infection: American Bullfrogs had the weakest association with infection while Western Toads had the strongest. After accounting for host species effects, hosts with greater snout-vent length had a lower probability of infection.

**Main conclusions:** Strong spatial influences at both biological levels suggest that mobile taxa (e.g., adult amphibians, birds, reptiles) may facilitate the movement of ranavirus among hosts and across the landscape. Higher taxonomic richness at sites may provide more opportunities for colonization or the presence of reservoir hosts that may influence ranavirus presence. Higher host richness correlating with higher ranavirus infection is suggestive of a dilution effect that has been observed for other amphibian disease systems and warrants further investigation. Our study demonstrates that an array of landscape, environmental, and host-level factors were associated with ranavirus epidemiology and illustrates that their importance vary with biological level.

## INTRODUCTION

Infectious diseases are increasingly recognized as important components of communities and ecosystems, yet their emergence in humans, wildlife, and plants across the globe has sparked concern because of their potentially devastating effects on populations (Daszak *et al.*, 2000; Dobson & Foufopoulos, 2001; Jones *et al.*, 2008). While decades of research have demonstrated the important roles of landscape and environmental (e.g., abiotic conditions and species interactions) processes in driving disease dynamics (reviewed in Poulin, 1998, 2007), a perpetual challenge in disease ecology is that the individual factors studied and their relative importance can be highly system-specific. For example, climate is cited as a major influence on vector-borne diseases (Githeko *et al.*, 2000; Rogers & Randolph, 2006; Rohr *et al.* 2011; Mordecai et al. 2017), flooding can influence the prevalence of cholera (reviewed in Ahern *et al.*, 2005), and loss of biodiversity can influence the prevalence of Lyme disease (Ostfeld & Keesing, 2000; Keesing *et al.*, 2006; Keesing *et al.*, 2010). Thus, for many emerging diseases, there is a need to conduct comprehensive field surveillance studies that combine assessments of key epidemiological parameters (e.g., presence, infection, pathogen load) with landscape and environmental data to determine the potential drivers of disease patterns across the landscape. Determining which factor—or groups of factors—is most influential can help to develop predictions, increase our knowledge base for host pathogen-interactions, and inform management and conservation (Rohr *et al.* 2015.

Recent studies have highlighted the importance of investigating the influence of factors at multiple biological levels of organization because of contrasting results between levels (e.g., site-versus individual-level; Borcard *et al.*, 2004; Dunn *et al.*, 2010; Schotthoefer *et al.*, 2011; Liu *et al.* 2013; Johnson *et al.*, 2015a; Cohen *et al.*, 2016). It has been hypothesized that abiotic factors influence distributional patterns at larger levels whereas biotic factors (e.g., species interactions) influence distributional patterns at smaller levels (Wiens, 1989; Levin, 1992; Rahbek, 2004; McGill, 2010; Cohen *et al.*, 2016). Accordingly, abiotic (e.g., temperature, precipitation, altitude) and biotic (e.g., host richness) factors were highly important in predicting the distribution of three pathogens (the pathogenic fungus *Batrachochytrium dendrobatidis* (Bd), West Nile virus, and the bacterium that causes Lyme disease (*Borrelia burgdorferi*), but biotic factors were more important at smaller levels (Cohen *et al.*, 2016). Landscape factors, such as connectivity among habitat patches, can also influence disease dynamics and the dispersal of pathogens. For example, the movement of the pathogenic fungus Bd through amphibian assemblages across the landscape suggests that dispersal plays a key role at regional levels (Laurance *et al.*, 1996; Lips *et al.*, 2008; Rohr *et al.* 2008; Vredenburg *et al.*, 2010; Liu *et al.* 2013). Therefore, evaluating which factors are most influential to the distribution of diseases, and at what levels of organization, is important to gain a clear understanding of what controls the spread of diseases among hosts *and* across the landscape.

Ranaviruses are viral pathogens of amphibians, fishes, and reptiles that have been implicated in mortality events across the globe (Duffus *et al.*, 2015). Over the last two decades, reports of mortality events in amphibian populations have gradually increased in the literature (Duffus *et al.*, 2015). Consequently, experimental studies and field surveys have been initiated to explore the potential drivers of ranavirus disease dynamics. Recent reviews have highlighted environmental factors that could influence ranaviral disease dynamics (Brunner *et al.*, 2015). For example, abiotic factors such as land use (e.g., cattle grazing and urbanization), water quality, and contaminants from runoff (e.g., nutrients, pesticides, heavy metals) are associated with increased prevalence of ranavirus in experimental studies and in the field (Forson & Storfer, 2006a; Forson & Storfer, 2006b; Kerby & Storfer, 2009; Kerby *et al.*, 2011; North *et al.*, 2015). In the United Kingdom (U.K.), deeper ponds were associated with an increased incidence of die-off events (North *et al.*, 2015). However, few studies have broadly explored the role of wetland characteristics on ranavirus occurrence or prevalence (Hoverman *et al.*, 2012a), particularly within an entire amphibian assemblage. In addition to abiotic factors, biotic factors (e.g., competition, predation, reservoir species) likely play a role in ranavirus distribution and dynamics. For instance, American Bullfrogs (*Rana catesbeiana*) and fish are implicated as potential reservoirs for the pathogen (Brunner *et al.*, 2015). It has also been hypothesized that predators can increase disease risk by inducing physiological stress that compromises immune function (Reeve *et al.*, 2013). Thus, while there are many hypothesized abiotic and biotic drivers of ranavirus emergence, there have been few attempts to assess the relative importance of these factors using large-scale field patterns for this pathogen.

The influences of landscape processes on ranavirus dynamics have received relatively little attention (Gahl & Calhoun, 2008; Hoverman *et al.*, 2012a; North *et al.*, 2015). Given that amphibians are often characterized by metapopulation dynamics (Gulve, 1994), the movement of infected hosts between breeding sites in close proximity to each other could influence spatial patterns in ranavirus occurrence on the landscape. Spatial models explained more variation than non-spatial models for ranavirus mortality events in the U.K. (North *et al.*, 2015; Price *et al.*, 2016). However, no spatial relationships were observed for mortality events in Acadia National Park, Maine, U.S.A (Gahl & Calhoun, 2008). An additional challenge is that most studies on the distribution of ranaviruses come from mortality events either detected by scientists or members of the public. This non-random selection of samples provides only sparse insight into the baseline epidemiology of ranaviruses in amphibian populations or the landscape and environmental processes underlying these patterns.

In the current study, our primary objective was to quantify the influence of a suite of landscape, abiotic, and biotic variables on ranavirus disease dynamics in amphibian assemblages. To this end, we conducted comprehensive field surveys of 93 wetlands to collect data on infection presence and prevalence within each amphibian population and obtain corresponding information on the biological and environmental characteristics associated with epidemiological observations. By collecting data from multiple amphibian host species and at both the individual and population (wetland) levels, we sought to broadly evaluate the influence of an array of factors on ranavirus epidemiology and how these factors influenced pathogen dynamics between two biological levels. To determine the relative influence of landscape, abiotic, and biotic factors on ranavirus, we used model selection and multi-model averaging followed by variance partitioning, thereby allowing us to assess the joint effects of hypothesized covariates and how they varied between the site-level and individual host-level.

## METHODS

### Study area and species

We examined patterns of ranavirus presence and infection in wetland amphibian assemblages located in the East Bay region of California (Figure 1; Hoverman *et al.*, 2012b; Johnson *et al.*, 2013b; Richgels *et al.*, 2013). We sampled 93 wetlands in managed parks and preserves within three counties (i.e., Alameda, Contra Costa, and Santa Clara; Johnson *et al.*, 2016). We selected wetlands that were smaller (< 2 ha) and likely to contain amphibian assemblages (Hoverman *et al.*, 2012b). Visitation to wetlands was haphazard, but was not spatiotemporally randomized because of logistical constraints. The amphibian assemblage in this region is composed of seven species: Northern Pacific Tree Frogs (*Hyliola regilla*), Western Toads (*Anaxyrus boreas*), American Bullfrogs (*R. catesbeiana*), California Newts (*Taricha torosa*), Rough-skinned Newts (*T. granulosa*), California Red-legged Frogs (*Rana draytonii*), and California Tiger Salamanders (*Ambystoma californiense*). Given the threatened status of California Red-legged Frogs and California Tiger Salamanders, we recorded them during surveys but excluded them from ranavirus sampling.

**Fig. 1.**
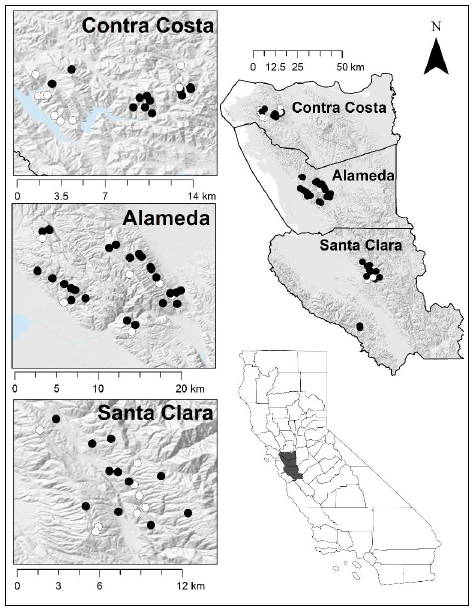
Study area and wetlands included in site-level analyses (*n* = 76) in three counties (Alameda, Contra Costa, and Santa Clara) of the East Bay region of California in 2013. Black points represent sites with ranavirus presence (those included in host-level analyses) and white points represent sites without ranavirus presence. Ranavirus was not detected at the southwest sites (those not included in the bottom inset map) within Santa Clara county.

### Field sampling, assessing ranavirus infection, and determining environmental variables

We conducted field surveys from May–August 2013 using the field sampling protocols of Hoverman et al. (2012b). In brief, we used a combination of visual encounter surveys, dipnet sweeps, and habitat-stratified seine hauls to sample the wetlands (Johnson *et al.*, 2013b; Richgels *et al.*, 2013). We disinfected all gear (e.g., nets and waders) with 15% bleach between sites. In the field, we identified amphibians to species, fishes to genus or species, and macroinvertebrates to order, family, or genus (Supplementary Table S1). At each wetland, we randomly selected up to 20 individuals (larvae, metamorphs, or both) per species for ranavirus screening. We necropsied individuals and sampled a portion of kidney and liver tissue for ranavirus. Equipment was flame sterilized between individuals. For each individual, ranaviral DNA was extracted from the combined liver and kidney tissue sample and infection was determined using standard quantitative PCR protocols (Forson & Storfer, 2006b).

We used an array of landscape, abiotic, and biotic predictor variables to represent environmental influences on ranavirus dynamics guided by theory and previous investigations (Table 1). Our landscape variable was distance to nearest ranavirus-infected wetland (other than the wetland the individual was found in). To calculate this distance, we recorded latitude and longitude of each site and measured Euclidean distance to nearest ranavirus-infected wetland using the R function ‘dist’. From the generated distance matrix, we deleted columns representing distances of each wetland to uninfected wetlands, and sorted to isolate distance to nearest ranavirus-infected wetland for each wetland and individual within each wetland. This method is limited in that not all wetlands in the landscape were sampled; thus, ranavirus-positive sites could occur, but not have been visited. However, our sampling scheme sought to sample all neighboring wetlands within a contiguous area (e.g., a park or preserve), such that these estimates are likely to capture general patterns related to colonization potential.

**Table 1.**
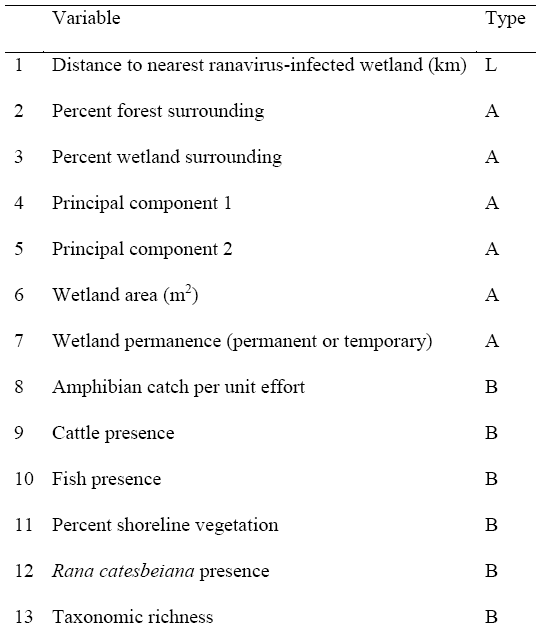
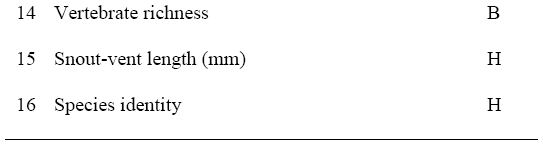
Predictor variables included to investigate patterns in landscape (L), abiotic (A), biotic (B), and host-level (H) influences on site-level ranavirus presence and host-level ranavirus infection in amphibian assemblages in the East Bay region of California in 2013. Host influences were only included in the host-level ranavirus infection analyses. Principal components 1 and 2 are the product of reducing the dimensionality of seven water quality parameters.

We assessed wetland permanence (permanent or temporary), percent forest or wetland surrounding wetlands, wetland area, and water quality factors at each site. We asses wetland permanence (permanent or temporary) based on water depth, wetland area, and with additional verification from historical images in Google Earth (Johnson *et al.*, 2013c). We measured conductivity (S/m), total dissolved solids (mg/l), salinity (mg/l), and pH with a YSI meter (Model 556; Yellow Spring Instrument, Yellow Springs, Ohio, USA). We quantified total nitrogen (mg/l), dissolved organic carbon (mg/l), and total ammonia (mg/l) using standard methods (http://snobear.colorado.edu/Kiowa/Kiowaref/procedure.html; Johnson *et al.*, 2013c). We used PCA to reduce dimensionality of the seven abiotic water-quality variables that we measured. Water-quality variables, except pH, were log-transformed to reduce positive skewness, and scaled and centered, before conducting the PCA. We retained only the first two components from PCA for further analyses, which had eigenvalues greater than one (Guttman-Kaiser criterion) and proportion of variance greater than the ‘broken-stick’ percentage (Supplementary Table S2; Yeomans & Golder, 1982; Kindt & Coe, 2005; Legendre & Legendre, 2012). Principal component 1 had high loadings for total dissolved solids (loading = −0.58), salinity (−0.57), and conductivity (−0.54). Principal component 2 was associated with total nitrogen (loading = 0.64), dissolved organic carbon (0.58), ammonium (0.46), and, to a lesser extent, pH (0.14). We calculated the percentage of area within a 1-km radius of each wetland classified as forested (sum of all forest types) and wetland (open water) using ArcGIS and the National Landcover Database (Johnson *et al.*, 2013b; Homer *et al.*, 2015) because of our interest in the influence of intact forest and wetlands surrounding focal wetlands. We calculated wetland surface area (m^2^; hereafter, area) by walking the perimeter of the pond with a handheld GPS using the track function. Area was base-10 log-transformed to meet assumptions of normality for analyses.

We represented the biotic community with percent vegetation cover on wetland shorelines (hereafter, percent shoreline vegetation), taxonomic richness, vertebrate richness, amphibian catch per unit effort (herein, CPUE), and the presence or absence of fishes, cattle, and non-native *R. catesbeiana*. We visually estimated percent shoreline vegetation at each site. We determined vertebrate richness by counting the number of amphibian and fish taxa. Taxonomic richness included all amphibians, fishes, and macroinvertebrates (detailed methods in Johnson *et al.*, 2016). We calculated CPUE by counting the number of individuals of each amphibian species during dip net sweeps and dividing by number of sweeps completed. We also included snout-vent length (mm), and species identity (*H. regilla, A. boreas*, *R. catesbeiana*, *T. torosa*, or *T. granulosa*) into host-level analyses. Snout-vent length was scaled and centered among species *Data analysis*

Our response variable for site-level analyses was ranavirus presence defined as one or more amphibians of any species infected with ranavirus within a wetland. We excluded wetlands with incomplete environmental data. Our response variable for host-level analyses was ranavirus infection defined as an individual having detectable ranavirus infection. We limited our ranavirus infection analyses only to wetlands where ranavirus was detected, which included infected and non-infected individuals. Therefore, we removed individuals from sites where ranavirus was not detected.

We assessed the influence of predictor variables on ranavirus presence and infection in amphibian assemblages with generalized linear models fitted with a binomial distribution and logit link. We conducted all analyses in program R v3.3.1 (R Development Core Team, 2015). We included base-10 log-transformed total number of individuals examined for ranavirus at each site as a fixed term to account for differences in the number of animals examined, which was expected to influence detection likelihood. For analyses of host-level infection, we used mixed effects models using the R package ‘lme4’ (Zuur *et al.*, 2009; Bates *et al.*, 2014) in which site was a random intercept term, thereby allowing us to nest observations from different amphibian species within the same site. We modeled host-level infection status (infected or not infected) to allow us to incorporate both host-level (e.g., body size) as well as site-level covariates (landscape, abiotic, and biotic). To keep models tractable, we initially used univariate analysis to identify associations between specific predictors variables and site-level ranavirus presence and host-level ranavirus infection. For univariate variable selection analyses, we used mixed model forms mentioned above (compared to correlations). Predictor variables with *P*-values < 0.10 from these univariate analyses were combined together into a global model. We centered and scaled all continuous predictor variables to facilitate comparison of coefficients among predictor variables and improve numerical stability. For snout-vent length of amphibian hosts, we centered and scaled within each species to account for differences in snout-vent length among species. We did not include interaction terms in global models because we did not hypothesize strong interactions between or among predictor variables, and to keep models tractable. We tested for collinearity between predictor variables included in the global models using Pearson’s correlation coefficients, and tested for multicollinearity among predictor variables in both global models with variance inflation factors with the R package ‘car’. We also calculated dispersion parameters to examine overdispersion in global models for ranavirus presence and prevalence. Additionally, we estimated the variance in site-level ranavirus presence and host-level ranavirus infection accounted for by landscape, abiotic, biotic, or individual variables in global models with the ‘varpart’ function in the R package ‘vegan’ (Borcard *et al.*, 1992; Schotthoefer *et al.*, 2011). We used the dredge function in the R package ‘MuMIn’ to create a set of all possible sub-models from ranavirus presence and infection global models and determine the best-supported model from the subset of predictor variables (Bartón, 2010). We compared models separately for ranavirus presence and infection analyses with an information-theoretic approach using Akaike’s Information Criterion (AIC; Burnham & Anderson, 2004; Mazerolle, 2016). We used AIC corrected for small sample sizes (AIC_C_) for both analyses because the number of observations divided by number of parameters was low for most ranavirus presence models (n/K < 40; Anderson & Burnham, 2002; Burnham & Anderson, 2004). Moreover, it is generally recommended to use AIC_C_ because it converges to AIC with large samples sizes such as we included in ranavirus infection analyses (Anderson & Burnham, 2002; Burnham & Anderson, 2004). We report model-averaged parameter estimates (β), standard errors (SE), adjusted SE, and relative importance of each predictor variable averaged from top models (ΔAIC_C_, < 4 AIC_C_ units). We investigated normality of response and predictor variables using kernel density plots and Q-Q plots, checked assumptions of all top models, and checked normality of model residuals against fitted values for top models. We investigated spatial autocorrelation of site-level ranavirus presence and residuals of ranavirus presence and infection global models using Moran’s I test in the R package ‘spdep’ (Borcard *et al.*, 1992; Schotthoefer *et al.*, 2011; Bivand, 2013).

## RESULTS

### Sampling overview

In total, our site-level analyses included 76 wetlands and 1,377 amphibians sampled for ranavirus representing five species. We removed 17 of the 93 originally surveyed sites from site-level analyses because they had incomplete site- or host-level covariate data, or both. The most common amphibian species among wetlands were *H. regilla* and *T. torosa*, and most sites (35%) had three amphibian species (Fig. 2). Thirty-two percent of tested amphibians were positive for ranavirus (*n* = 441 of 1,377). At least one infected individual occurred at 67% of wetlands (*n* = 51 of 76) and an average of 61% of individuals were infected at wetlands with ranavirus infection (95% CI = 53–68%). For host-level analyses, we removed 288 individuals from 25 sites where ranavirus was not present; thus, we reduced our host-level sample size to 1,089 individuals. The percentage of infected individuals at wetlands varied among species; *T. granulosa* had the highest average percentage of individuals infected (mean = 60%, 95% CI = 48–71%) followed by *A. boreas* (36%, 26–45%), *T. torosa* (25%, 20–30%), *H. regilla* (25%, 20–30%), and *R. catesbeiana* (16%, 6–25%). We observed non-native *R. catesbeiana* at 29% (*n* = 22) of wetlands and fish presence (i.e., *Gambusia affinis*, *Lepomis macrochirus*, *Carassius auratus*, *Ictalurus* spp., or *Micropterus* spp.) was observed at 26% of wetlands (*n* = 20).

**Fig. 2.**
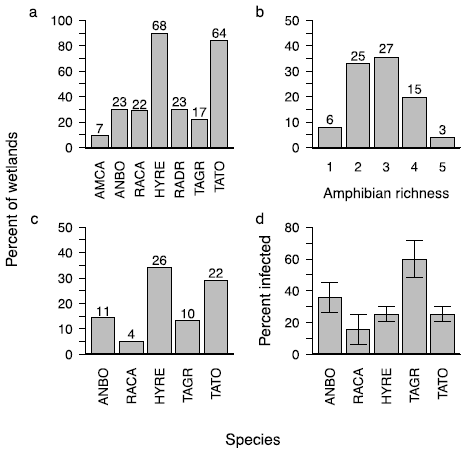
Percent of wetlands with each species (a), species richness at wetlands (b), percent of wetlands with ranavirus infected hosts for each species (c), and mean percent of hosts infected with ranavirus per wetland (with 95% CI) of those collected of each species (d) in amphibian assemblages in the East Bay region of California in 2013. Numbers above bars indicate number of wetlands with each species or species richness (*n* = 76). For plots (a), (c), and (d): AMCA, *Ambystoma californiense* (California Tiger Salamander); ANBO, *Anaxyrus boreas* (Western Toad); RACA, *Rana catesbeiana* (American Bullfrog); HYRE, *Hyliola regilla* (Northern Pacific Tree Frog); RADR, *Rana draytonii* (California Red-legged Frog); TAGR, *Taricha granulosa* (Rough-skinned Newt); TATO, *Taricha torosa* (California Newt).

### Model selection and multimodel inference

Univariate analyses determined that landscape (distance to nearest ranavirus-infected wetland), abiotic (percent wetland), and biotic (CPUE and taxonomic richness) were associated with, and included in the global model for, site-level ranavirus presence. For host-level ranavirus infection, univariate analyses demonstrated that landscape (distance to nearest ranavirus-infected wetland), abiotic (percent wetland), biotic (*R. catesbeiana* presence and vertebrate richness), and host-level (snout-vent length and species identity) were associated with, and included in the global model for, host-level ranavirus infection. From the global models, we produced 16 total models comprised of four landscape and abiotic variables for site-level analysis of ranavirus presence and 64 total models comprised of six landscape, abiotic, biotic, and host-level variables for host-level analysis of ranavirus infection using the dredge function in R (Supplementary material Appendix 1, Tables A3 and A4). For site-level ranavirus presence analysis, four models were within 4 AIC_C_ of the best-supported model (Supplementary material Appendix 1, Table A5). For host-level ranavirus infection analysis, eight models were within 4 AIC_C_ of the best-supported model (Supplementary material Appendix 1, Table A6).

Landscape and biotic variables had the strongest associations with site-level ranavirus presence in our best-supported models (Table 2). Distance to nearest ranavirus-infected wetland and taxonomic richness were included in all best-supported models, while CPUE and nearby wetland area were only included half of the best supported-models. Wetlands that were farther from the nearest ranavirus-infected wetland had a lower likelihood of ranavirus presence (β = −0.26 ± 0.05 [model-averaged coefficient ± adjusted SE]; Fig. 3). Wetlands with greater taxonomic richness had a higher likelihood of ranavirus presence (β = 0.12 ± 0.04). Variance partitioning analyses demonstrated that the landscape variable, distance to nearest ranavirus-infected wetland, explained the most variance (adjusted R^2^ from variance partitioning = 0.18) and the biotic variable, taxonomic richness, explained a smaller portion of variance (R^2^ = 0.09) in site-level ranavirus presence (Table 3).

**Table 2.** Model-averaged coefficients for centered and scaled predictor variables from a subset of models (delta AICc < 4 points, 4 of 16 models) of site-level ranavirus presence in amphibian assemblages in the East Bay region of California in 2013. Coefficients are arranged by ascending *P*-value, then alphabetically. “Distance” is distance to nearest ranavirus-infected wetland (km), “Num. mod.” is the number of models that include that predictor variable, “Importance” is percent of models within the model subset that contain that variable, “SE” is standard error, and “Adj. SE” is adjusted standard error. Coefficients with *P* ≤ 0.05 are shaded in grey.

| Variable | Num. mod. | Importance | Estimate | SE | Adj. SE | *z* | *P* |
| --- | --- | --- | --- | --- | --- | --- | --- |
| Distance | 4 | 1.00 | -0.26 | 0.05 | 0.05 | 5.24 | < 0.001 |
| Taxonomic richness | 4 | 1.00 | 0.12 | 0.04 | 0.04 | 2.61 | < 0.001 |
| Catch per unit effort | 2 | 0.57 | 0.09 | 0.05 | 0.05 | 1.65 | 0.099 |
| Total dissected | 4 | 1.00 | 0.03 | 0.06 | 0.06 | 0.59 | 0.554 |
| Percent wetland | 2 | 0.23 | 0.01 | 0.06 | 0.06 | 0.10 | 0.919 |

**Table 3.**
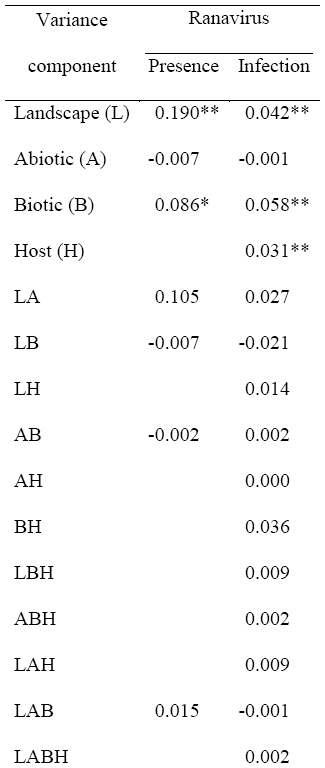
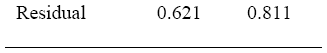
Results of variance partitioning analyses quantifying the amount of unique variation (adjusted R^2^) to landscape, abiotic, biotic, and host-level (Host) variables, and the shared variation between and among the variable subsets, for site-level ranavirus presence and host-level ranavirus infection. Host-level variables were only included in host-level analyses and probability values can only be calculated for landscape, abiotic, biotic, and host-level components. An asterisk (*) indicates *P* < 0.01 and two asterisks (**) indicate P < 0.001

| Variance<br>component | Ranavirus |  |
| --- | --- | --- |
|  | Presence | Infection |
| Landscape (L) | 0.190** | 0.042** |
| Abiotic (A) | -0.007 | -0.001 |
| Biotic (B) | 0.086* | 0.058** |
| Host (H) |  | 0.031** |
| LA | 0.105 | 0.027 |
| LB | -0.007 | -0.021 |
| LH |  | 0.014 |
| AB | -0.002 | 0.002 |
| AH |  | 0.000 |
| BH |  | 0.036 |
| LBH |  | 0.009 |
| ABH |  | 0.002 |
| LAH |  | 0.009 |
| LAB | 0.015 | -0.001 |
| LABH |  | 0.002 |
| Residual | 0.621 | 0.811 |

**Fig. 3.**
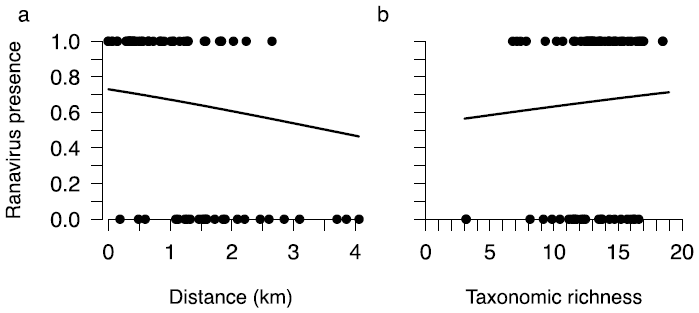
Model-averaged predicted probability of site-level ranavirus presence (*n* = 76) in amphibian assemblages in the East Bay region of California in 2013 with increasing (a) distance to nearest ranavirus-infected wetland (Distance, km), and (b) taxonomic richness in wetlands in 2013. Points for taxonomic richness are jittered to reduce overlap.

The best-supported models for host-level ranavirus infection prevalence included landscape, abiotic, biotic, and host-level predictor variables (Table 4). Distance to nearest ranavirus-infected wetland, snout-vent length, species identity, and vertebrate richness had the strongest associations with ranavirus infection. Hosts in wetlands that were further from the nearest ranavirus-infected wetland had the lowest likelihood of ranavirus infection (distance β = −1.40 ± 0.38; Fig. 4). Hosts in wetlands with greater vertebrate richness, while controlling for host density, were less likely to be infected (β = −0.61 ± 0.31). Additionally, species differed in their likelihood of ranavirus infection. *Rana catesbeiana*, which was the reference level in the species identity variable, had the lowest likelihood of ranavirus infection (β = −4.09 ± 0.90; Fig. 5). *Taricha torosa* (β = 2.66 ± 0.84), *P. regilla* (β = 3.02 ± 0.84), *A. boreas* (β = 3.72 ± 0.85), and *T. granulosa* (β = 4.03 ± 0.92) had higher likelihood of ranavirus infection relative to *R. catesbeiana*. Finally, hosts with greater snout-vent length were less likely to be infected (β = −0.40 ± 0.11). Variance partitioning demonstrated that the biotic variable, taxonomic richness explained the most variation in ranavirus infection at the host-level (adjusted R^2^ = 0.06) followed by landscape (adjusted R^2^ = 0.04) and host-level variables (species identity and snout-vent length; adjusted R^2^ = 0.03; Table 3).

**Table 4.**
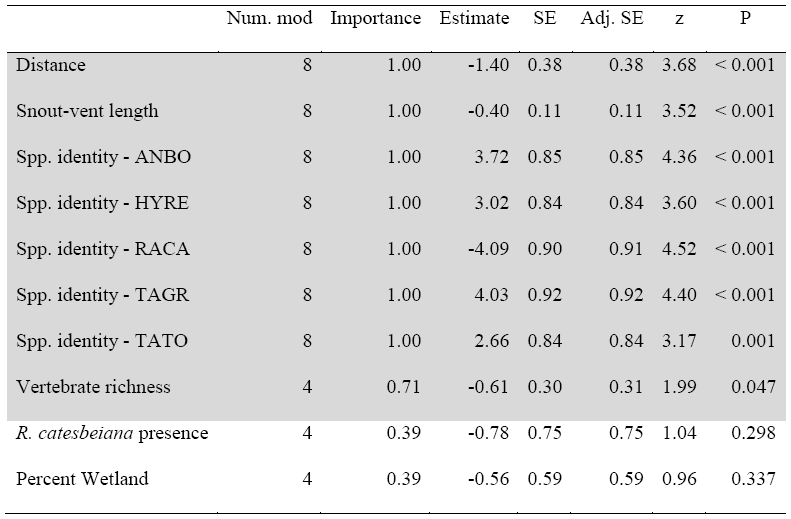
Model-averaged coefficients for centered and scaled predictor variables from a subset of models (delta AICc < 4 points, 8 of 64 models) of host-level ranavirus infection in amphibian assemblages in the East Bay region of California in 2013. Coefficients are arranged by ascending *P*-value, then alphabetically. “Distance” is distance to nearest ranavirus-infected, “Num. mod.” is the number of models that include that predictor variable, “Importance” is percent of models within the model subset that contain that variable, “SE” is standard error, and “Adj. SE” is adjusted standard error. For species identity (Spp. identity), “ANBO” is *A. boreas*, “HYRE” is *H. regilla*, “RACA” is *R. catesbeiana* (the reference level), “TAGR” is *T. granulosa*, and “TATO” is *T. torosa*. Coefficients with *P* ≤ 0.05 are shaded in grey.

|  | Num. mod | Importance | Estimate | SE | Adj. SE | z | P |
| --- | --- | --- | --- | --- | --- | --- | --- |
| Distance | 8 | 1.00 | -1.40 | 0.38 | 0.38 | 3.68 | < 0.001 |
| Snout-vent length | 8 | 1.00 | -0.40 | 0.11 | 0.11 | 3.52 | < 0.001 |
| Spp. identity - ANBO | 8 | 1.00 | 3.72 | 0.85 | 0.85 | 4.36 | < 0.001 |
| Spp. identity - HYRE | 8 | 1.00 | 3.02 | 0.84 | 0.84 | 3.60 | < 0.001 |
| Spp. identity - RACA | 8 | 1.00 | -4.09 | 0.90 | 0.91 | 4.52 | < 0.001 |
| Spp. identity - TAGR | 8 | 1.00 | 4.03 | 0.92 | 0.92 | 4.40 | < 0.001 |
| Spp. identity - TATO | 8 | 1.00 | 2.66 | 0.84 | 0.84 | 3.17 | 0.001 |
| Vertebrate richness | 4 | 0.71 | -0.61 | 0.30 | 0.31 | 1.99 | 0.047 |
| *R. catesbeiana* presence | 4 | 0.39 | -0.78 | 0.75 | 0.75 | 1.04 | 0.298 |
| Percent Wetland | 4 | 0.39 | -0.56 | 0.59 | 0.59 | 0.96 | 0.337 |

**Fig. 4.**
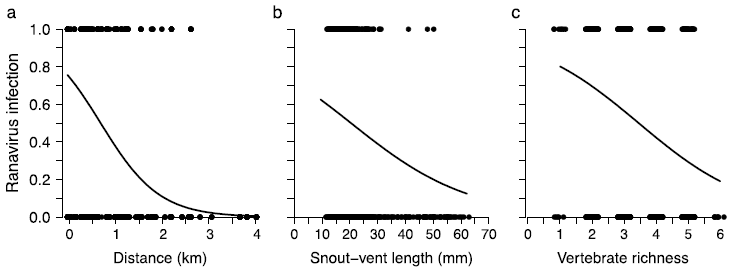
Model-averaged predicted probability of host-level ranavirus infection (*n* = 1,089) in amphibian assemblages in the East Bay region of California in 2013 with increasing (a) distance from nearest ranavirus-infected wetland (Distance, km), (b) snout-vent length, and (c) vertebrate richness. Points for vertebrate richness are jittered to reduce overlap.

**Fig. 5.**
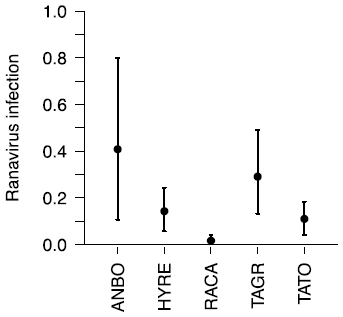
Model-averaged predicted probability of ranavirus infection with standard error (*n* = 1,089) for amphibian hosts in the East Bay region of California in 2013. For species: ANBO, *Anaxyrus boreas* (Western Toad); HYRE, *Hyliola regilla* (Northern Pacific Tree frog); RACA, *Rana catesbeiana* (American Bullfrog; the reference level); TAGR, *Taricha granulosa* (Rough-skinned Newt); and TATO, *Taricha torosa* (California Newt).

No spatial autocorrelation was observed for ranavirus presence (*P* = 0.865) in site-level observations based on Moran’s I. Additionally, residuals for ranavirus presence and infection models with the most support were not spatially autocorrelated based on Moran’s I (*P* > 0.792). Collinearity between predictor variables was low; however, and as expected, collinearity was highest between distance to nearest ranavirus-infected wetland and the amount of nearby wetland area in both analyses (ρ = 0.64 and 0.61). Variance inflation factors (VIFs) for all predictor variables in ranavirus presence and infection global models indicated low multicollinearity among variables (VIFs < 2.27). Overdispersion was not observed in site-level ranavirus presence and host-level infection global models (dispersion parameters < 1).

## DISCUSSION

For any infectious disease, it is critical to identify the landscape and environmental factors that influence the distribution of the pathogen to develop a broader understanding of disease emergence and strategies for management and conservation. Here, we examined the landscape and environmental factors underlying patterns in site-level ranavirus presence and host-level ranavirus infection in amphibian assemblages in the East Bay region of California during 2013. We used comprehensive field surveillance data, rather than observations of mortality events commonly used to describe patterns in ranavirus disease dynamics, and model selection with multimodel inference to determine ranavirus epidemiology in the assemblage. Ranavirus was widespread throughout our study site and our analyses demonstrated that site- and host-level patterns in ranavirus epidemiology were more strongly associated with landscape and biotic factors (aspects of species richness), rather than abiotic factors.

At the landscape level, wetlands in closer proximity to ranavirus-positive wetlands were more likely to support ranavirus and have higher infection prevalence. To date, the influence of landscape processes on ranavirus dynamics is poorly understood. Disease risk might be greatest for wetlands in close proximity to other infected wetlands, which has been found in other amphibian disease systems. For example, the movement of the pathogenic fungus Bd through amphibian assemblages across the landscape suggests that dispersal probably plays an important role (Laurance *et al.*, 1996; Lips *et al.*, 2008; Rohr *et al.* 2008; Vredenburg *et al.*, 2010; Liu *et al.* 2013). Previous research has found equivocal results related to the spatial clustering of ranavirus-associated mortality events (Gahl & Calhoun, 2008; North *et al.*, 2015). Our findings suggest that the movement of infected amphibians among wetlands could distribute ranavirus from infected wetlands to other nearby wetlands. Amphibians can metamorphose from wetlands with ranavirus infections and the returning adults can harbor infections (Brunner *et al.*, 2004). For instance, a reconstructed ranavirus emergence event in the U.K. demonstrated a localized spread from nearby ponds with distances spread similar to known amphibian and frog dispersal distances (Price *et al.*, 2016). While this suggests that infected hosts can move ranavirus across the landscape, the movement patterns of infected hosts have not been explored. Given that the dispersal ability of most amphibians is relatively limited (Blaustein *et al.*, 1994; Wells, 2010), the probability of infected hosts reaching distant wetlands is relatively low. In our study, there was a 20 and 60% reduction in ranavirus presence and infection, respectively, at about 2 km.

Wetlands in close proximity to ranavirus-positive wetlands might have more frequent introductions of the virus into the system thereby increasing exposure and infection probabilities. Movement of other taxa (e.g., reptiles, birds, humans) either via sublethally infected hosts or immune taxa transporting ranavirus on their surfaces could also distribute ranavirus across the landscape (reviewed in Brunner *et al.*, 2015). However, the transfer of ranavirus on the surface of immune taxa might be rare given that ranaviruses can be rapidly degraded in the environment by naturally occurring plankton and microbes (Johnson & Brunner, 2014) and when wetland drying occurs (Brunner *et al.*, 2007). Ranavirus could also be distributed across the landscape when rain events and flooding occur, which can connect nearby wetlands through the movement of water. Future research examining the movement of ranavirus-infected hosts and other sources of ranavirus dispersal among wetlands will provide critical information on how ranavirus moves across the landscape and influences disease risk.

The influence of biodiversity on disease risk has been a major focus of recent disease ecology research (Keesing *et al.*, 2006; Civitello *et al.* 2015; Johnson *et al.*, 2015b). Although rarely considered in ranavirus studies, we found that biotic factors broadly related to species richness were associated with ranavirus patterns. In our study, taxonomic richness correlated positively with the probability of ranavirus presence at the site-level whereas vertebrate richness was correlated negatively with host-level ranavirus infection prevalence. Greater taxonomic richness could increase the likelihood that ranavirus is introduced into a wetland (e.g., via mobile taxa) or the probability of successfully establishing in a species, as also found in other studies of parasites (e.g., Johnson *et al.*, 2013a; Rottstock *et al.*, 2014; Johnson *et al.*, 2016). Additionally, more diverse wetlands might support potential reservoirs for ranavirus infection, although there was no evidence that fish or non-native Bullfrog were associated with patterns in ranavirus infection. The negative association between vertebrate richness and infection is suggestive of a dilution effect, yet our field data lack estimates of transmission within the communities to confirm this mechanism. The dilution effect has been observed in other amphibian disease systems (trematodes and B. dendrobatidis; Searle *et al.*, 2011; Johnson *et al.*, 2013a; Venesky *et al.*, 2014a,b; Rohr *et al.*, 2015) and therefore might also occur for ranavirus. Because this is the first study to document associations between species richness and ranavirus dynamics, the mechanisms underlying these patterns are in need of further investigation with controlled experiments.

Although environmental stressors have frequently been hypothesized as drivers of ranavirus epidemiology (Gray *et al.*, 2007; Greer & Collins, 2008; Brunner *et al.*, 2015), we found no significant interactions between ranavirus occurrence and the factors representing environmental stressors that we measured in this study. For instance, factors associated with cattle (i.e. cattle presence, reduced shoreline vegetation, increased ammonia) did not influence ranavirus presence or infection in our analyses. Additionally, there was no association with the amount of forest surrounding the wetlands, which functions as an indicator of habitat integrity. Lastly, there was no evidence that non-native *R. catesbeiana* or fishes contributed to ranavirus patterns, despite the postulated importance of these groups as reservoirs of ranavirus and other amphibian pathogens in other regions (Brunner *et al.*, 2015).

Host-level factors such as amphibian species identity were also a major factor in explaining infection prevalence. *Rana catesbeiana* exhibited the lowest likelihood of infection among the five species sampled in these wetlands. *Rana catesbeiana* had only 3% overall infection prevalence, even after accounting for site-level differences. Using *R. catesbeiana* as the references species, infection tended to be higher in the remaining species. Our findings are similar to previous laboratory experiments where *R. catesbeiana* were relatively resistant to ranavirus infection compared to other amphibian species (Hoverman *et al.*, 2011). For the remaining species in the assemblage, there is a need to conduct experimental studies examining their susceptibility to ranavirus. Preliminary results from our research group have found high levels of susceptibility to infection and high mortality in *H. regilla*, and moderate infection and mortality in *A. boreas* (N.M. Hambalek, personal communication).

We observed that larger host body size (greater snout-vent length) reduced the probability of ranavirus infection, even after accounting for species-level differences in body size. This observation coincides with an observation that body size was negatively associated with Bd infection or Bd-induced death (Rohr *et al.* 2013; Gervasi *et al.*, 2017), but positive (Raffel *et al.* 2013; McMahon *et al.* 2014) and non-linear relationship (Raffel *et al.* 2010) have also been observed. It also coincides with frequent observations that juveniles might be more prone to infection than adults (i.e., with larger body sizes) in amphibians and fishes (Cullen *et al.*, 1995; Ariel & Owens, 1997; Cullen & Owens, 2002; Jensen *et al.*, 2011). Larger body size may be an indicator of a more-developed immune system, which could prevent infections from establishing (Miller *et al.*, 2011; Gervasi *et al.*, 2017). Future field-based and experimental studies investigating relationships among size, development, and ranavirus infection will undoubtedly benefit our understanding of ranavirus infection in amphibians.

## CONCLUSIONS

Despite decades of research on ranavirus-amphibian interactions, our understanding of the factors underlying ranavirus epidemiology in natural systems remains limited. While numerous factors have been proposed as drivers of infection, it still remains unclear why the outcome of a ranavirus outbreak can vary from no obvious mortality to a massive die-off event (Brunner et al. 2015). Moreover, the predominant focus on ranavirus-associated mortality events has failed to capture baseline epidemiological patterns across the landscape. Using a dataset from 76 wetlands, five amphibian species, and 1,377 hosts, our results illustrate that landscape and biotic factors were most important for explaining ranavirus epidemiology. In particular, landscape factors explained more variance at larger (site-level) biological levels while biotic and host-level factors explained more variance at smaller biological levels (host-level). Our findings are similar to those suggested for other disease distributions and highlight the importance of investigating factors influencing disease epidemiology at multiple biological levels (Schotthoefer *et al.*, 2011; Johnson *et al.*, 2015a; Cohen *et al.*, 2016). Several variables such as cattle presence and water chemistry parameters, that are often cited to influence ranavirus epidemiology (Forson & Storfer, 2006a; Forson & Storfer, 2006b; Kerby & Storfer, 2009; Kerby *et al.*, 2011), were not influential in our study. Additionally, the variables we included in our analyses explained scant variability in site-level ranavirus presence and host-level ranavirus infection. Therefore, further experimental and field-based investigations of proposed and novel factors will undoubtedly help broaden our understanding of the dynamics of this emerging infectious pathogen and benefit management and conservation.

## ACKNOWLEDGMENTS

We thank Melina Allahverdian, Kelly DeRolf, Jackie Gregory, Emily Hannon, Jeremy Henderson, Megan Housman, Aaron Klingborg, Bryan LaFonte, Keegan McCaffrey, Mary Toothman, and Vanessa Wuerthner for assistance with data collection and sample processing. We also thank Michael Chislock, Erin Kenison, Emily McCallen, and Katherine Richgels for helpful discussions of data analyses. This research was supported by funding from the National Institutes of Health, Ecology, and Evolution of Infectious Diseases Program grant (R01GM109499), the National Science Foundation (DEB 1149308), and the David and Lucile Packard Foundation. For access to properties and logistical support, we thank the East Bay Regional Parks District, the East Bay Municipal Utility District, Santa Clara County Parks, Blue Oak Ranch Reserve (specific thanks to Michael Hamilton), California State Parks, The Nature Conservancy, and many private landowners. The content is solely the responsibility of the authors and does not necessarily represent the official views of the National Institutes of Health, Ecology, and Evolution of Infectious Diseases.

## SUPPORTING INFORMATION

**Table S1.** Sampled amphibian, fish, and macroinvertebrate species in the East Bay Region of California in 2013.

**Table S2.** Results of principal component analysis for seven abiotic water variables and ‘broken-stick’ test.

**Table S3.** Results of univariate variable selection for correlation between site-level ranavirus presence and 14 predictor variables.

**Table S4.** Results of univariate variable selection for correlation between host-level ranavirus infection and 16 predictor variables.

**Table S5.** Summary statistics for the four top-ranked models for site-level ranavirus presence.

**Table S6.** Summary statistics for the eight top-ranked models for host-level ranavirus presence.

## DATA ACCESSIBILITY STATEMENT

Data used for spatial analyses and mapping available from the National Land Cover Database (https://catalog.data.gov/dataset/national-land-cover-database-nlcd-land-cover-collection) and the State of California Geoportal (http://portal.gis.ca.gov/geoportal). Ranavirus infection and microhabitat data included in this study are available from the Pangea database (data deposited, will insert link when posted and after review).

## BIOSKETCH

**Brian Tornabene** is a PhD Student at the University of Montana in Missoula, Montana, USA. His work focuses on natural history, ecotoxicology, and population and disease ecology—often with herpetofauna. Currently, he is investigating the influence of brine contamination from oil and gas development on amphibian communities.

## Literature Cited

Ahern, M., Kovats, R.S., Wilkinson, P., Few, R. & Matthies, F. (2005). Global health impacts of floods: epidemiologic evidence. Epidemiologic reviews, 27, 36–46.

Anderson, D.R. & Burnham, K.P. (2002). Avoiding pitfalls when using information-theoretic methods. Journal of Wildlife Management, 66, 912–918.

Ariel, E. & Owens, L. (1997). Epizootic mortalities in tilapia Oreochromis mossambicus. Diseases of Aquatic Organisms, 29, 1–6.

Bartón, K. (2010). MuMIn: multi-model inference, 2010. R package version, 1

Bates, D., Mächler, M., Bolker, B. & Walker, S. (2014). Fitting linear mixed-effects models using lme4. arXiv preprint arXiv:1406.5823,

Bivand, R.S. (2013). spdep: Spatial Dependence: Weighting Schemes, Statistics and Models.

Blaustein, A.R., Wake, D.B. & Sousa, W.P. (1994). Amphibian declines: judging stability, persistence, and susceptibility of populations to local and global extinctions. Conservation Biology, 8, 60–71.

Borcard, D., Legendre, P. & Drapeau, P. (1992). Partialling out the Spatial Component of Ecological Variation. Ecology, 73, 1045–1055.

Borcard, D., Legendre, P., Avois-Jacquet, C. & Tuomisto, H. (2004). Dissecting the spatial structure of ecological data at multiple scales. Ecology, 85, 1826–1832.

Brunner, J.L., Schock, D.M. & Collins, J.P. (2007). Transmission dynamics of the amphibian ranavirus Ambystoma tigrinum virus. Dis Aquat Organ, 77, 87-95.

Brunner, J.L., Schock, D.M., Davidson, E.W. & Collins, J.P. (2004). Intraspecific reservoirs: Complex life history and the persistence of a lethal ranavirus. Ecology, 85, 560–566.

Brunner, J.L., Storfer, A., Gray, M.J. & Hoverman, J.T. (2015). Ranavirus ecology and evolution: From epidemiology to extinction. Ranaviruses: Lethal pathogens of ectothermic vertebrates (ed. by M.J. Gray and G.D. Chinchar), pp. 71–104. Springer, New York, U.S.A.

Burnham, K.P. & Anderson, D.R. (2004). Multimodel inference - understanding AIC and BIC in model selection. Sociological Methods & Research, 33, 261–304.

Civitello, D.J., Cohen, J., Fatima, H., Halstead, N.T., Liriano, J., McMahon, T.A., Ortega, C.N., Sauer, E.L., Sehgal, T., Young, S. & Rohr, J.R. (2015) Biodiversity inhibits parasites: Broad evidence for the dilution effect. Proceedings of the National Academy of Sciences of the United States of America, 112, 8667-8671.

Cohen, J.M., Civitello, D.J., Brace, A.J., Feichtinger, E.M., Ortega, C.N., Richardson, J.C., Sauer, E.L., Liu, X. & Rohr, J.R. (2016). Spatial scale modulates the strength of ecological processes driving disease distributions. Proceedings of the National Academy of Science of the United States of America, 113, E3359–64.

Cullen, B. & Owens, L. (2002). Experimental challenge and clinical cases of Bohle iridovirus (BIV) in native Australian anurans. Diseases of aquatic organisms, 49, 83–92.

Cullen, B., Owens, L. & Whittington, R. (1995). Experimental infection of Australian anurans (Limnodynastes terraereginae and Litoria latopalmata) with Bohle iridovirus. Diseases of Aquatic Organisms, 23, 83–92.

Daszak, P., Cunningham, A.A. & Hyatt, A.D. (2000). Emerging infectious diseases of wildlife—threats to biodiversity and human health. Science, 287, 443.

Dobson, A. & Foufopoulos, J. (2001). Emerging infectious pathogens of wildlife. Philosophical Transactions of the Royal Society of London Series B-Biological Sciences, 356, 1001–1012.

Duffus, A.L.J., Waltzek, T.B., Stöhr, A.C., Allender, M.C., Gotesman, M., Whittington, R.J., Hick, P., HInes, M.K. & Marschang, R. (2015). Distribution and Host Range of Ranaviruses. Ranaviruses: Lethal pathogens of ectothermic vertebrates (ed. by M.J. Gray and G.D. Chinchar), pp. 9–57. Springer, New York, U.S.A.

Dunn, R.R., Davies, T.J., Harris, N.C. & Gavin, M.C. (2010). Global drivers of human pathogen richness and prevalence. Proceedings of the Royal Society of London B: Biological Sciences, 277, 2587–95.

Forson, D. & Storfer, A. (2006a). Effects of atrazine and iridovirus infection on survival and life-history traits of the long-toed salamander (*Ambystoma macrodactylum*). Environmental Toxicology and Chemistry, 25, 168–173.

Forson, D.D. & Storfer, A. (2006b). Atrazine increases ranavirus susceptibility in the tiger salamander, *Ambystoma tigrinum*. Ecological Applications, 16, 2325–2332.

Gahl, M.K. & Calhoun, A.J.K. (2008). Landscape setting and risk of *Ranavirus* mortality events. Biological Conservation, 141, 2679–2689.

Gervasi, S.S., Stephens, P.R., Hua, J., Searle, C.L., Xie, G.Y., Urbina, J., Olson, D.H., Bancroft, B.A., Weis, V., Hammond, J.I., Relyea, R.A. & Blaustein, A.R. (2017). Linking Ecology and Epidemiology to Understand Predictors of Multi-Host Responses to an Emerging Pathogen, the Amphibian Chytrid Fungus. PLoS One, 12, e0167882.

Githeko, A.K., Lindsay, S.W., Confalonieri, U.E. & Patz, J.A. (2000). Climate change and vector-borne diseases: a regional analysis. Bulletin of the World Health Organization, 78, 1136–1147.

Gray, M.J., Miller, D.L., Schmutzer, A.C. & Baldwin, C.A. (2007). *Frog virus* 3 prevalence in tadpole populations inhabiting cattle-access and non-access wetlands in Tennessee, USA. Diseases of Aquatic Organisms, 77, 97–103.

Greer, A.L. & Collins, J.P. (2008). Habitat fragmentation as a result of biotic and abiotic factors controls pathogen transmission throughout a host population. Journal of Animal Ecology, 77, 364–369.

Gulve, P.S. (1994). Distribution and extinction patterns within a northern metapopulation of the pool frog, *Rana lessonae*. Ecology, 75, 1357–1367.

Homer, C., Dewitz, J., Yang, L.M., Jin, S., Danielson, P., Xian, G., Coulston, J., Herold, N., Wickham, J. & Megown, K. (2015). Completion of the 2011 National Land Cover Database for the Conterminous United States - Representing a Decade of Land Cover Change Information. Photogrammetric Engineering and Remote Sensing, 81, 345–354.

Hoverman, J.T., Gray, M.J., Haislip, N.A. & Miller, D.L. (2011). Phylogeny, life history, and ecology contribute to differences in amphibian susceptibility to ranaviruses. Ecohealth, 8, 301–319.

Hoverman, J.T., Gray, M.J., Miller, D.L. & Haislip, N.A. (2012a). Widespread occurrence of ranavirus in pond-breeding amphibian populations. Ecohealth, 9, 36–48.

Hoverman, J.T., Mihaljevic, J.R., Richgels, K.L.D., Kerby, J.L. & Johnson, P. (2012b). Widespread co-occurrence of virulent pathogens within california amphibian communities. Ecohealth, 9, 288–292.

Jensen, B.B., Holopainen, R., Tapiovaara, H. & Ariel, E. (2011). Susceptibility of pike-perch Sander lucioperca to a panel of ranavirus isolates. Aquaculture, 313, 24–30.

Johnson, A.F. & Brunner, J.L. (2014). Persistence of an amphibian ranavirus in aquatic communities. Diseases of Aquatic Organisms, 111, 129–138.

Johnson, P.T., de Roode, J.C. & Fenton, A. (2015a). Why infectious disease research needs community ecology. Science, 349, 1259504.

Johnson, P.T., Wood, C.L., Joseph, M.B., Preston, D.L., Haas, S.E. & Springer, Y.P. (2016). Habitat heterogeneity drives the host-diversity-begets-parasite-diversity relationship: evidence from experimental and field studies. Ecology Letters, 19, 752–61.

Johnson, P.T.J., Ostfeld, R.S. & Keesing, F. (2015b). Frontiers in research on biodiversity and disease. Ecology Letters, 18, 1119–1133.

Johnson, P.T.J., Preston, D.L., Hoverman, J.T. & LaFonte, B.E. (2013a). Host and parasite diversity jointly control disease risk in complex communities. Proceedings of the National Academy of Sciences of the United States of America, 110, 16916–16921.

Johnson, P.T.J., Preston, D.L., Hoverman, J.T. & Richgels, K.L.D. (2013b). Biodiversity decreases disease through predictable changes in host community competence. Nature, 494, 230–233.

Johnson, P.T.J., Hoverman, J.T., McKenzie, V.J., Blaustein, A.R. & Richgels, K.L.D. (2013c). Urbanization and wetland communities: applying metacommunity theory to understand the local and landscape effects. Journal of Applied Ecology, 50, 34–42.

Jones, K.E., Patel, N.G., Levy, M.A., Storeygard, A., Balk, D., Gittleman, J.L. & Daszak, P. (2008). Global trends in emerging infectious diseases. Nature, 451, 990–994.

Keesing, F., Holt, R.D. & Ostfeld, R.S. (2006). Effects of species diversity on disease risk. Ecology Letters, 9, 485–498.

Keesing, F., Belden, L.K., Daszak, P., Dobson, A., Harvell, C.D., Holt, R.D., Hudson, P., Jolles, A., Jones, K.E. & Mitchell, C.E. (2010). Impacts of biodiversity on the emergence and transmission of infectious diseases. Nature, 468, 647–652.

Kerby, J.L. & Storfer, A. (2009). Combined effects of atrazine and chlorpyrifos on susceptibility of the tiger salamander to *Ambystoma tigrinum* virus. Ecohealth, 6, 91–98.

Kerby, J.L., Hart, A.J. & Storfer, A. (2011). Combined effects of virus, pesticide, and predator cue on the larval tiger salamander (*Ambystoma tigrinum*). Ecohealth, 8, 46–54.

Kindt, R. & Coe, R. (2005). Tree diversity analysis: A manual and software for common statistical methods for ecological and biodiversity studies. World Agroforestry Centre.

Laurance, W.F., McDonald, K.R. & Speare, R. (1996). Epidemic disease and the catastrophic decline of Australian rain forest frogs. Conservation Biology, 10, 406–413.

Legendre, P. & Legendre, L. (2012). Numerical ecology, Third English edition. edn. Elsevier, Amsterdam.

Levin, S.A. (1992). The problem of pattern and scale in ecology: the Robert H. MacArthur award lecture. Ecology, 73, 1943–1967.

Lips, K.R., Diffendorfer, J., Mendelson, J.R. & Sears, M.W. (2008). Riding the wave: Reconciling the roles of disease and climate change in amphibian declines. PloS Biology, 6, 441–454.

Liu, X., Rohr, J.R. & Li, Y. (2013) Climate, vegetation, introduced hosts and trade shape a global wildlife pandemic. Proceedings of the Royal Society of London B: Biological Sciences, 280, 20122506.

Mazerolle, M.J. (2016). Model Selection and Multimodel Inference Based on (Q)AIC(c). Documentation for R: A language and environment for statistical computing. R Foundation for Statistical Computing.

McGill, B.J. (2010). Ecology. Matters of scale. Science, 328, 575–6.

McMahon, T.A., Sears, B.F., Venesky, M.D., Bessler, S.M., Brown, J.M., Deutsch, K., Halstead, N.T., Lentz, G., Tenouri, N., Young, S., Civitello, D.J., Ortega, N., Fites, J.S., Reinert, L.K., Rollins-Smith, L.A., Raffel, T.R. & Rohr, J.R. (2014) Amphibians acquire resistance to live and dead fungus overcoming fungal immunosuppression. Nature, 511, 224-227.

Miller, D.L., Gray, M.J. & Storfer, A. (2011). Ecopathology of ranaviruses infecting amphibians. Viruses, 3, 2351–2373.

Mordecai, E.A., Cohen, J.M., Evans, M.V., Gudapati, P., Johnson, L.R., Lippi, C.A., Miazgowicz, K., Murdock, C.C., Rohr, J.R. & Ryan, S.J. (2017) Detecting the impact of temperature on transmission of Zika, dengue, and chikungunya using mechanistic models. PLoS Neglected Tropical Diseases, 11, e0005568.

North, A.C., Hodgson, D.J., Price, S.J. & Griffiths, A.G.F. (2015). Anthropogenic and Ecological Drivers of Amphibian Disease (Ranavirosis). Plos One, 10, e0127037.

Ostfeld, R.S. & Keesing, F. (2000). Biodiversity and disease risk: the case of Lyme disease. Conservation Biology, 14, 722–728.

Poulin, R. (1998). Evolutionary ecology of parasites: from individuals to communities. Chapman & Hall, New York, New York, U.S.A.

Poulin, R. (2007). The Evolutionary Ecology of Parasites, 2nd edn. Princeton University Press, Princton, NJ.

Price, S.J., Garner, T.W., Cunningham, A.A., Langton, T.E. & Nichols, R.A. (2016). Reconstructing the emergence of a lethal infectious disease of wildlife supports a key role for spread through translocations by humans. Proceedings of the Royal Society B, 283, 20160952.

R Development Core Team (2015). R: A language and environment for statistical computing. R Foundation for Statistical Computing, Vienna, Austria. ISBN 3-900051-07-0, URL http://www.r-project.org/.

Raffel, T.R., Halstead, N.T., McMahon, T., Romansic, J.M., Venesky, M.D. & Rohr, J.R. (2013) Disease and thermal acclimation in a more variable and unpredictable climate. Nature Climate Change, 3, 146-151.

Raffel, T.R., Michel, P.J., Sites, E.W. & Rohr, J.R. (2010) What drives chytrid infections in newt populations? Associations with substrate, temperature, and shade. EcoHealth, 7, 526-536.

Rahbek, C. (2004). The role of spatial scale and the perception of large-scale species-richness patterns. Ecology Letters, 8, 224–239.

Reeve, B.C., Crespi, E.J., Whipps, C.M. & Brunner, J.L. (2013). Natural stressors and ranavirus susceptibility in larval wood frogs (Rana sylvatica). EcoHealth, 10, 190–200.

Richgels, K.L.D., Hoverman, J.T. & Johnson, P.T.J. (2013). Evaluating the role of regional and local processes in structuring a larval trematode metacommunity of Helisoma trivolvis. Ecography, 36, 854–863.

Rogers, D. & Randolph, S. (2006). Climate change and vector-borne diseases. Advances in parasitology, 62, 345–381.

Rohr, J.R., Civitello, D.J., Crumrine, P.W., Halstead, N.T., Miller, A.D., Schotthoefer, A.M., Stenoien, C., Johnson, L.B. & Beasley, V.R. (2015). Predator diversity, intraguild predation, and indirect effects drive parasite transmission. Proceedings of the National Academy of Sciences, 112, 3008–3013.

Rohr, J.R., Dobson, A.P., Johnson, P.T.J., Kilpatrick, A.M., Paull, S.H., Raffel, T.R., Ruiz-Moreno, D. & Thomas, M.B. (2011) Frontiers in climate change-disease research. Trends in Ecology & Evolution, 26, 270-277.

Rohr, J.R., Raffel, T.R., Blaustein, A.R., Johnson, P.T.J., Paull, S.H. & Young, S. (2013) Using physiology to understand climate-driven changes in disease and their implications for conservation. Conservation Physiology, 1, doi:10.1093/conphys/cot022.

Rohr, J.R., Raffel, T.R., Halstead, N.T., McMahon, T.A., Johnson, S.A., Boughton, R.K. & Martin, L.B. (2013) Early-life exposure to a herbicide has enduring effects on pathogen-induced mortality. Proceedings of the Royal Society B-Biological Sciences, 280, 20131502.

Rohr, J.R., Raffel, T.R., Romansic, J.M., McCallum, H. & Hudson, P.J. (2008) Evaluating the links between climate, disease spread, and amphibian declines. Proceedings of the National Academy of Sciences of the United States of America, 105, 17436-17441.

Rottstock, T., Joshi, J., Kummer, V. & Fischer, M. (2014). Higher plant diversity promotes higher diversity of fungal pathogens, while it decreases pathogen infection per plant. Ecology, 95, 1907-1917.

Schotthoefer, A.M., Rohr, J.R., Cole, R.A., Koehler, A.V., Johnson, C.M., Johnson, L.B. & Beasley, V.R. (2011). Effects of wetland vs. landscape variables on parasite communities of Rana pipiens: links to anthropogenic factors. Ecological Applications, 21, 1257–1271.

Searle, C.L., Biga, L.M., Spatafora, J.W. & Blaustein, A.R. (2011). A dilution effect in the emerging amphibian pathogen *Batrachochytrium dendrobatidis*. Proceedings of the National Academy of Sciences of the United States of America, 108, 16322–16326.

Venesky, M.D., Liu, X., Sauer, E.L. & Rohr, J.R. (2014a). Linking manipulative experiments to field data to test the dilution effect. Journal of Animal Ecology, 83, 557–65.

Venesky, M.D., Raffel, T.R., McMahon, T.A. & Rohr, J.R. (2014b) Confronting inconsistencies in the amphibian-chytridiomycosis system: implications for disease management. Biological Reviews, 89, 477-483.

Vredenburg, V.T., Knapp, R.A., Tunstall, T.S. & Briggs, C.J. (2010). Dynamics of an emerging disease drive large-scale amphibian population extinctions. Proceedings of the National Academy of Sciences of the United States of America, 107, 9689–9694.

Wells, K.D. (2010). The ecology and behavior of amphibians. University of Chicago Press.

Wiens, J.A. (1989). Spatial scaling in ecology. Functional ecology, 3, 385–397.

Yeomans, K.A. & Golder, P.A. (1982). The guttman-kaiser criterion as a predictor of the number of common factors. Journal of the Royal Statistical Society Series D-the Statistician, 31, 221–229.

Zuur, A.F., Leno, E.N., Wlaker, N., Saveliev, A.A. & Smith, G.M. (2009). Mixed effects models and extensions in ecology with R. Springer, New York, NY.

